# Evidence for quinol oxidase activity of ImoA, a novel NapC/NirT family protein from the neutrophilic Fe(II) oxidizing bacterium *Sideroxydans lithotrophicus* ES-1

**DOI:** 10.1101/2022.01.10.475773

**Authors:** Abhiney Jain, Anaísa Coelho, Joana Madjarov, Smilja Todorovic, Ricardo O. Louro, Jeffrey A. Gralnick, Catarina M. Paquete

## Abstract

The freshwater chemolithoautotrophic Gram-negative bacterium *Sideroxydans lithotrophicus* ES-1 oxidizes Fe(II) at the cell surface. In this organism, it is proposed that the monoheme cytochrome MtoD from the Mto pathway transfer electrons across the periplasm to an inner membrane NapC/NirT family tetraheme cytochrome encoded by *Slit_2495*, for which we propose the name ImoA (<u>i</u>nner <u>m</u>embrane <u>o</u>xidoreductase). ImoA has been proposed to function as the quinone reductase, receiving electrons from iron oxidizing extracellular electron uptake pathway to reduce the quinone pool. In this study, ImoA was cloned on a pBAD plasmid vector and overexpressed in *Escherichia coli*. Biochemical and spectroscopic characterization of the purified ImoA reveals that this 26.5 kDa cytochrome contains one high-spin and three low-spin hemes. Our data show that ImoA can function as a quinol oxidase and is able to functionally replace CymA, a related NapC/NirT family tetraheme cytochrome required for anaerobic respiration of a wide range of substrates by *Shewanella oneidensis*. We demonstrate that ImoA can transfer electrons to different periplasmic proteins from *S. oneidensis* including STC and FccA, but in a manner that is distinct from that of CymA. Phylogenetic analysis shows that ImoA is clustered closer to NirT sequences than to CymA. This study suggests that ImoA functions as a quinol oxidase in *S. oneidensis* and raises questions about the directionality and/or reversibility of electron flow through the Mto pathway in *S. lithotrophicus* ES-1.

**Importance:** Fe(II)-oxidizing bacteria play an important role in the biogeochemical cycling of iron, representing a promising class of organisms for the development of novel biotechnological processes, including bioelectrosynthesis. These organisms perform extracellular electron transfer, taking up electrons from Fe(II) outside of the cell, possibly through a porin-cytochrome complex in the outer membrane. The electrons are then transferred to the quinone pool in the inner membrane via periplasmic and inner membrane electron transfer proteins. In this paper, we produced and characterized the NapC/NirT family tetraheme cytochrome ImoA, encoded by *Slit_2495*, an inner membrane protein from the Gram-negative Fe(II)-oxidizing bacterium *Sideroxydans lithotrophicus* ES-1, proposed to be involved in extracellular electron transfer to the quinone pool. We show that ImoA may function instead as a quinol oxidase. The obtained insights represent the first step in understanding mechanisms of electron flow in *S. lithotrophicus* ES-1 and may lead towards practical biotechnological applications of Fe(II)-oxidizing bacteria.

## Introduction

The contribution of chemolithoautotrophic Fe(II)-oxidizing bacteria (FeOB) to iron cycling in the environment has been widely recognized in recent years (1). These organisms rely on Fe(II) as the electron donor and energy source while reducing oxygen to fix carbon dioxide (1–3). FeOB play an important role in the biogeochemical cycling of iron, as well as in the development of novel biotechnological processes, including bioelectrosynthesis (4). Despite the relevance of these bacteria, the functionality of electron transfer proteins required for their growth and survival in the environment is primarily based on genomic analysis, heterologous expression and biochemical analysis (5–8). Although a genetic system has been reported for the FeOB bacterium *Mariprofundus ferrooxydans* (9), methods to generate gene deletions are still lacking, which are critical tools for validating predictions of functionality.

The oxidation of Fe(II) to Fe(III) under circumneutral pH conditions in oxic environments forms insoluble iron oxyhydroxides. Therefore, FeOB evolved strategies to prevent intracellular Fe(III) precipitation, including catalyzing Fe(II) oxidation at the cellular surface (6). These organisms perform extracellular electron transfer, taking up electrons from Fe(II) outside of the cell using cell-surface or periplasmic proteins that have access to extracellular environment through an outer membrane porin to oxidize Fe(II) (2, 10, 11). The electrons released from Fe(II) oxidation are then thought to be transferred to quinone pools in the inner membrane via periplasmic and inner membrane electron transfer proteins, typically *c*-type cytochromes (5).

The freshwater chemolithoautotrophic Gram-negative bacterium *Sideroxydans lithotrophicus* ES-1 (12), has been proposed to oxidize Fe(II) at the cell surface through the decaheme porin-cytochrome complex MtoAB (6). MtoA and MtoB are homologs of MtrA and MtrB from the MtrCAB complex from the Fe(III) reducing Gram-negative bacterium *Shewanella oneidensis* MR-1, known to be involved in outward extracellular electron transfer (13). *S. oneidensis* MR-1 is a metabolically versatile facultative anaerobic bacterium able to reduce a large number of soluble and insoluble electron acceptors (14–16). Extracellular electron transfer by the Mtr pathway requires a host of multiheme cytochromes including CymA, STC, FccA, MtrA, and MtrC. These cytochromes create a conductive pathway to transfer electrons from the inner membrane, across the periplasm and the outer-membrane for the reduction of both soluble and solid electron acceptors outside of the cell (17–20).

The *mtoA* and *mtoB* genes are located in a cluster encoded by *Slit2495-2498* on the *S. lithotrophicus* ES-1 genome. This cluster also contains two other *c*-type cytochromes: a putative periplasmic monoheme cytochrome MtoD that is proposed to transfer electrons across the periplasm, and a NapC/NirT family tetraheme cytochrome encoded by *Slit_2495* (21). Slit_2495 was proposed to function as the quinone reductase during Fe(II) oxidation based on its sequence similarity with the tetraheme cytochrome CymA from *S. oneidensis* MR-1 (6). In *S. oneidensis* MR-1, CymA acts as an electron hub that receives electrons from the quinone pool, transferring the electrons to numerous periplasmic proteins (17, 22). All sequenced chemolithoautotrophic FeOB, including *S. lithotrophicus* ES-1, are predicted to encode a *bc*_1_ complex, which is proposed to reduce the quinone pool required to generate NADH (5, 23). The presence of a *bc*_1_ complex (23) and similarity of Slit_2495 to CymA, a quinone oxidoreductase (21), makes the possible quinone reductase role of Slit_2495 protein redundant and speculative.

To understand the possible functionality of this protein, we heterologously expressed Slit_2495 and characterized its biochemical and electrochemical properties. These insights, together with phylogenetic analysis and *in vivo* assays where CymA was replaced by Slit_2495 in *S. oneidensis*, reveal that Slit_2495 may function as a quinol oxidase in *S. lithotrophicus* ES-1. Slit_2495 has been putatively called CymA in the literature (5, 6). However, phylogenetic analysis shows that Slit_2495 is more closely related to NirT from *Pseudomonas* than to CymA from *Shewanella*, and our data demonstrate that although it may function as a quinol oxidase as CymA, the respective electron transfer mechanisms to periplasmic proteins in *Shewanella* are different. Based on our characterization, we propose that Slit_2495 be named ImoA for <u>i</u>nner <u>m</u>embrane <u>o</u>xidoreductase.

## Results

### Phylogenetic analysis of ImoA

Phylogenetic analysis of ImoA was performed with respect to other NapC/NirT family proteins. We included sequences of CymA from *Shewanella*, NirT from *Pseudomonas*, NrfH from *Desulfovibrio* and cytochrome *c*552 sequences from *Nitrosomonas europaea* along with NapC/NirT family protein sequences from *Sideroxydans*. Sequences from TorC or DorC were not included in the analysis since both of these proteins contain five heme binding sites compared to four heme binding sites in ImoA. Our phylogenetic analysis shows that ImoA clustered close to NapC sequences from *Escherichia coli* and NirT sequences from *Pseudomonas* (**Figure 1**). Furthermore, the protein sequence of ImoA is 64% identical to NirT from *Pseudomonas stutzeri* and 54% identical to NapC from *E. coli*, whereas it is only 33% identical to CymA from *S. oneidensis* MR-1.

**Figure 1.**
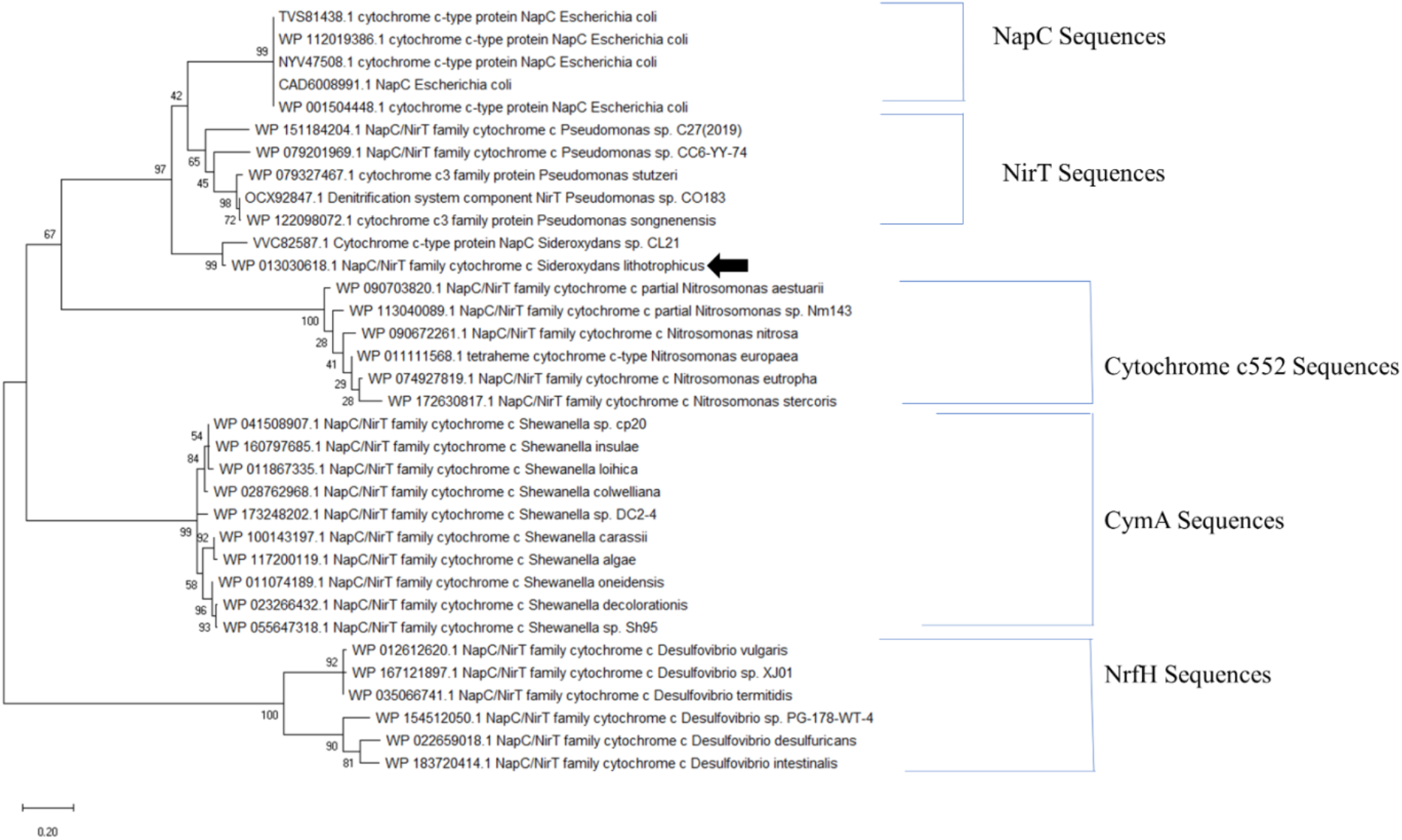
Phylogenetic analysis of ImoA with respect to other NapC/NirT proteins. Maximum likelihood tree was built using 2,000 bootstrap replications. Clusters of different NapC/NirT protein families used in the analysis are identified by the brackets labelled with specific proteins and black arrow points to ImoA sequence. The tree is drawn to scale, and the scale bar represents substitutions per site.

### Production of ImoA

Biochemical and electrochemical properties of ImoA were characterized employing purified protein. To facilitate purification the protein was engineered to carry a C-terminal Strep-II affinity tag (**Figure S1**). After purification, SDS-PAGE analysis reveals that ImoA was pure and ran as a single band with an apparent molecular weight between 20 and 25 kDa (**Figure S2A**). The band stained positively for covalently attached hemes (**Figure S2B**), and N-terminal sequencing retrieved the predicted sequence of ImoA (amino acid sequence: NNKTG). Mass spectrometry revealed that the purified ImoA has a molecular mass of ~26.5 kDa, which agrees with the calculated molecular mass of the apo-protein (22.9 kDa) with the incorporation of the 4 hemes (~0.6 kDa per heme) and the Strep-tag (1 kDa). Pure ImoA shows an absorbance ratio of A_Soret Peak_/A_280nm_ of 3.5 (**Figure 2A**).

**Figure 2.**
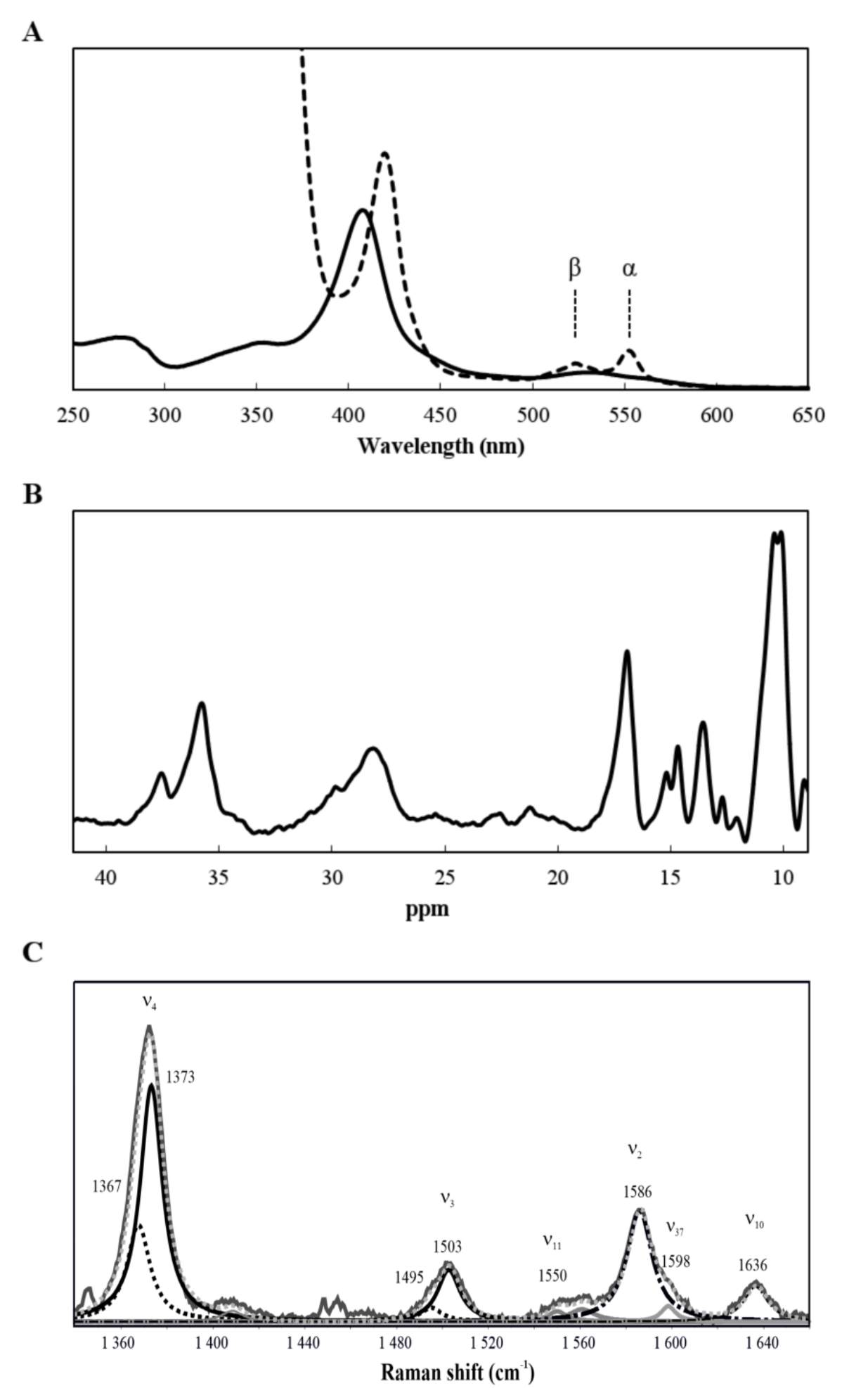
Spectroscopic properties of ImoA. (A) UV-visible spectroscopy in the oxidized (solid line) and reduced state (dashed line); (B) ^1^H-1D-NMR spectrum obtained in the oxidized state; (C) High frequency region of RR spectra. Experimental spectrum (light grey, solid line) and overall deconvoluted spectrum (light grey, dotted line), with designated ν_4_ and ν_3_ high-spin (black dotted trace) and low-spin (black solid trace) bands, non-assigned and spin state nonspecific bands (grey solid trace) and spin-state specific, non-deconvoluted bands ν_2_ and ν_10_ (black dot-dashed trace).

Gel filtration was performed to provide direct insights into the oligomeric state of ImoA in the conditions used in the present study. The n-Dodecyl β-D-maltoside (DDM)-solubilized ImoA was eluted with an apparent molecular mass of approximately 80 kDa. Given that the mass of a DDM micelle is approximately 51 kDa (24), and of ImoA approximately 26 kDa, the results obtained indicate that ImoA is a monomer within the DDM micelle.

### ImoA contains low- and high-spin hemes

UV-visible spectra of ImoA show the typical features of a low-spin *c*-type cytochrome, with the characteristic Soret peak in the oxidized protein with a maximum at 408 nm (**Figure 2A**). Upon reduction of ImoA, the Soret peak shifts to 420 nm, and the α and β peaks appear with maxima at 552 and 523 nm, respectively. In a low-spin cytochrome, the Nuclear Magnetic Resonance (NMR) signals of the methyl groups of the hemes in the oxidized state of the protein are shifted to the paramagnetic region of the spectrum (17, 25). The ^1^H-1D-NMR spectrum of ImoA in the oxidized state exhibits the typical features of a cytochrome with low-spin hemes, with signals appearing from 15 ppm to 40 ppm, being well resolved from the protein envelope region (**Figure 2B**). The presence of low-spin *c*-type cytochrome in the ^1^H-1D-NMR spectrum of ImoA is consistent with the results obtained by UV-visible spectroscopy. In the next step, we addressed the spin state of the hemes by employing highly sensitive Resonance Raman (RR) spectroscopy, which can unambiguously detect different spin/coordination states in multiple heme containing proteins. High frequency region of RR spectra of ImoA reveals heme vibrational marker bands (v_i_) that are asymmetric and broadened. The most sensitive indicators of the heme spin and redox state, ν_4_ and ν_3_ bands, suggest the presence of two ferric heme populations. Their identity becomes evident upon component analysis of the spectra, which allowed the determination of the exact frequencies of these bands. The ν_4_ at 1373 cm^−1^ and ν_3_ at 1503 cm^−1^ are characteristic of low-spin state, which is predominant, while ν_4_ at 1367 cm^−1^ and ν_3_ at 1495 cm^−1^ suggest the presence of a high-spin population (**Figure 2C**) (26, 27). The relative ratio of the band intensities (i.e. ν_3_ (LS) / ν_3_ (HS)), together with the fact that the protein contains 4 hemes, indicate the presence of one high-spin and three low-spin hemes.

### Redox properties of ImoA

To evaluate the redox properties of ImoA, pure protein was adsorbed to the surface of a pyrolytic graphite edge (PGE) electrode, and cyclic voltammetry was performed (**Figure 3**). ImoA titrates between 0 and −400 mV. Although in a protein with multiple redox centers, the voltammogram contain information about the redox properties of each individual heme, the discrimination of these values is not straightforward due to the interaction between the redox and ionizable centers (28). Indeed, it is possible to observe differences in the voltammograms for the different pH values (**Figure S3**), suggesting that the reduction potentials of the individual hemes are different and changes with the pH.

**Figure 3.**
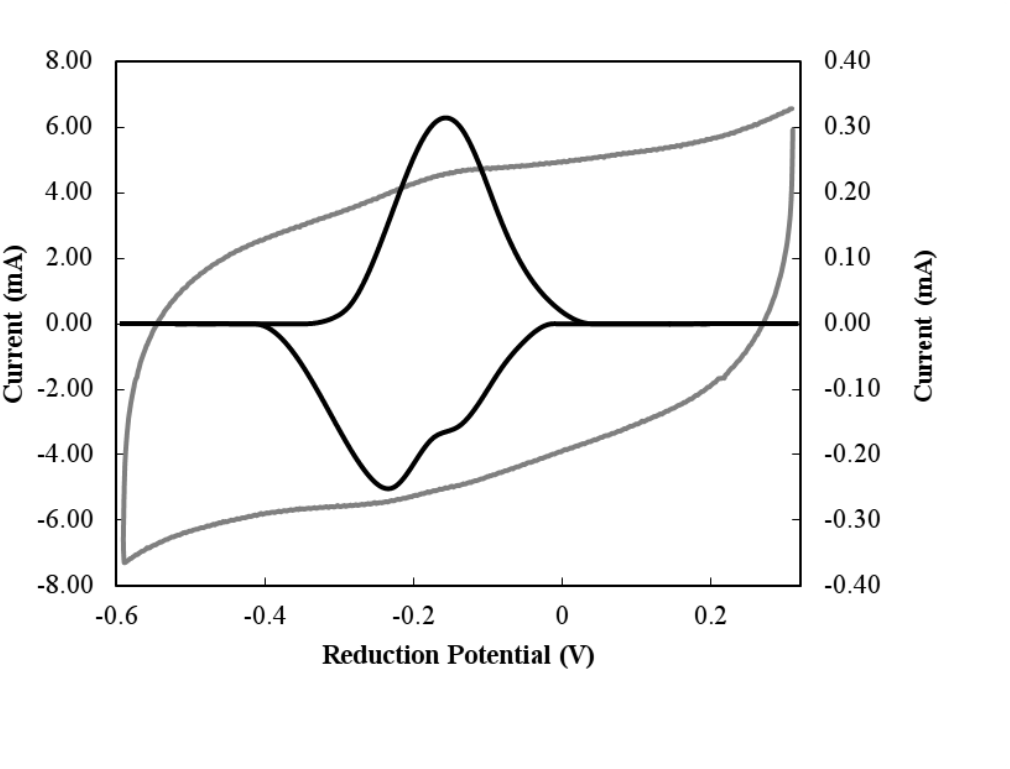
Cyclic voltammetry of ImoA. Raw (black and left axis) and baseline-subtracted data (grey and right axis) of the voltammogram obtained at a scan rate of 400 mV/s at pH 7.8.

### ImoA can function as a quinol oxidase in the Mtr pathway of *S. oneidensis*

The quinol oxidase activity of ImoA was investigated by expressing and testing it in a *S. oneidensis* deletion mutant lacking CymA (Δ*cymA*) (**Figure 4**). ImoA is able to replace CymA and complement the ferric citrate reduction activity in Δ*cymA* (**Figure 4A**). The Δ*cymA* strain containing pBBR1MCS-2::*imoA* reduced ferric citrate to the same extent as a Δ*cymA* strain complemented with pBBR1MCS-2::*cymA*, although at a slightly slower rate. *In vivo* experiments of Δ*cymA* strains containing pBBR1MCS-2::*cymA*, pBBR1MCS-2::*imoA* or an empty pBBR1MCS-2 vector for approximately 40 h show their ability to perform extracellular electron transfer to anodes (**Figure 4B**). While Δ*cymA* carrying an empty plasmid produced negligible current, the two Δ*cymA* strains harboring either the native *cymA* or *imoA* produced approximately 20 μA/cm^2^ of geometric anode surface, demonstrating that ImoA takes up the role of CymA in *S. oneidensis* for electron transport to an anode. These results show that ImoA functions as a quinol oxidase and transfers electrons to periplasmic proteins in *S. oneidensis*.

**Figure 4.**
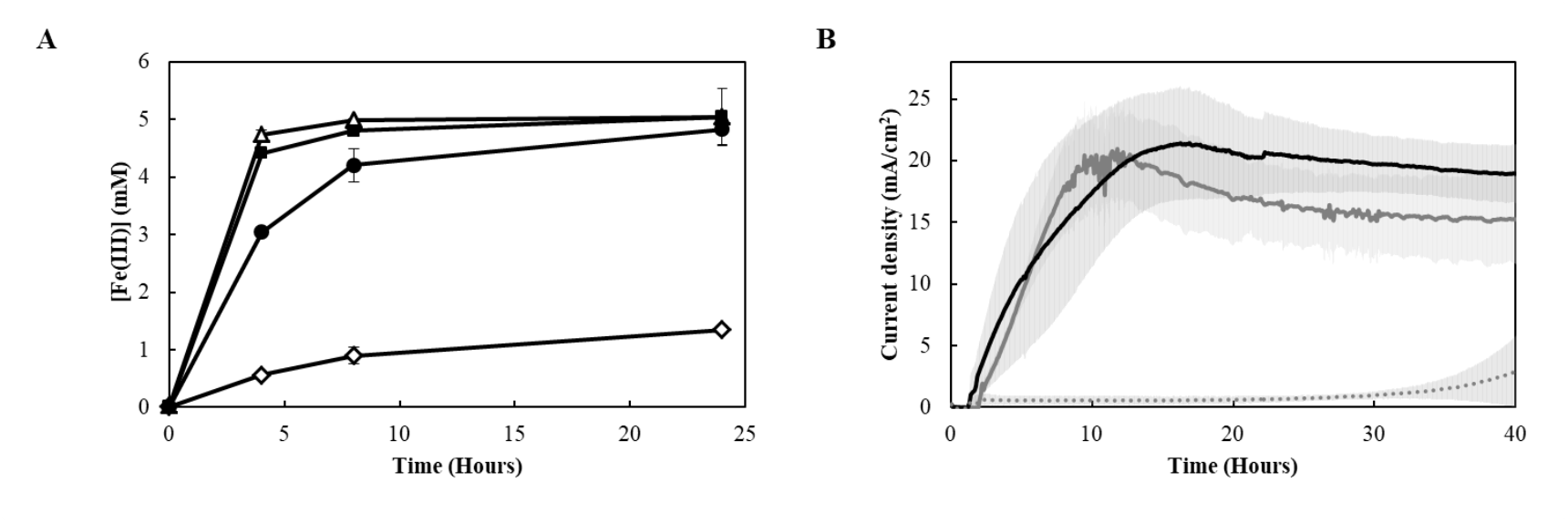
Quinol oxidase functionality of ImoA in the Mtr pathway. **(A)** Resting cell ferric citrate reduction: *S. oneidensis* containing pBBR1MCS-2 (Δ), Δ*cymA* containing pBBR1MCS-2 (◊), Δ*cymA* containing pBBR1MCS-2::*cymA* (■) and Δ*cymA* containing pBBR1MCS-2::*imoA* (●). Error bars represent standard deviation of the mean from experiments performed in triplicate. (B) Current density produced by the different Δ*cymA* strains containing pBBR1MCS-2:: *imoA* (black line), pBBR1MCS-2:: *cymA* (grey line), or empty pBBR1MCS-2 (grey dotted line) in a bioelectrochemical reactor. Error bars shown in gray represent standard deviation of the mean from experiments performed in two triplicates.

### ImoA exchanges electrons with multiple periplasmic proteins

Given that CymA is able to exchange electrons with multiple periplasmic proteins in *S. oneidensis* (17, 22), we tested the ability of Δ*cymA* expressing ImoA to grow on different electron acceptors that require CymA, including fumarate (**Figure 5A**), dimethyl sulfoxide (DMSO) (**Figure 5B**) and nitrate (**Figure 5C**). ImoA is able to complement the growth of Δ*cymA* strains with the electron acceptors tested, however the growth of the strains containing pBBR1MCS-2:: *imoA* is slower than those containing pBBR1MCS-2:: *cymA*. This indicates that, although ImoA can replace CymA, the electron transfer between the quinol oxidase and the different periplasmic proteins appears less efficient. No growth was observed on any of the electron acceptors for Δ*cymA* containing an empty pBBR1MCS-2 vector.

**Figure 5.**
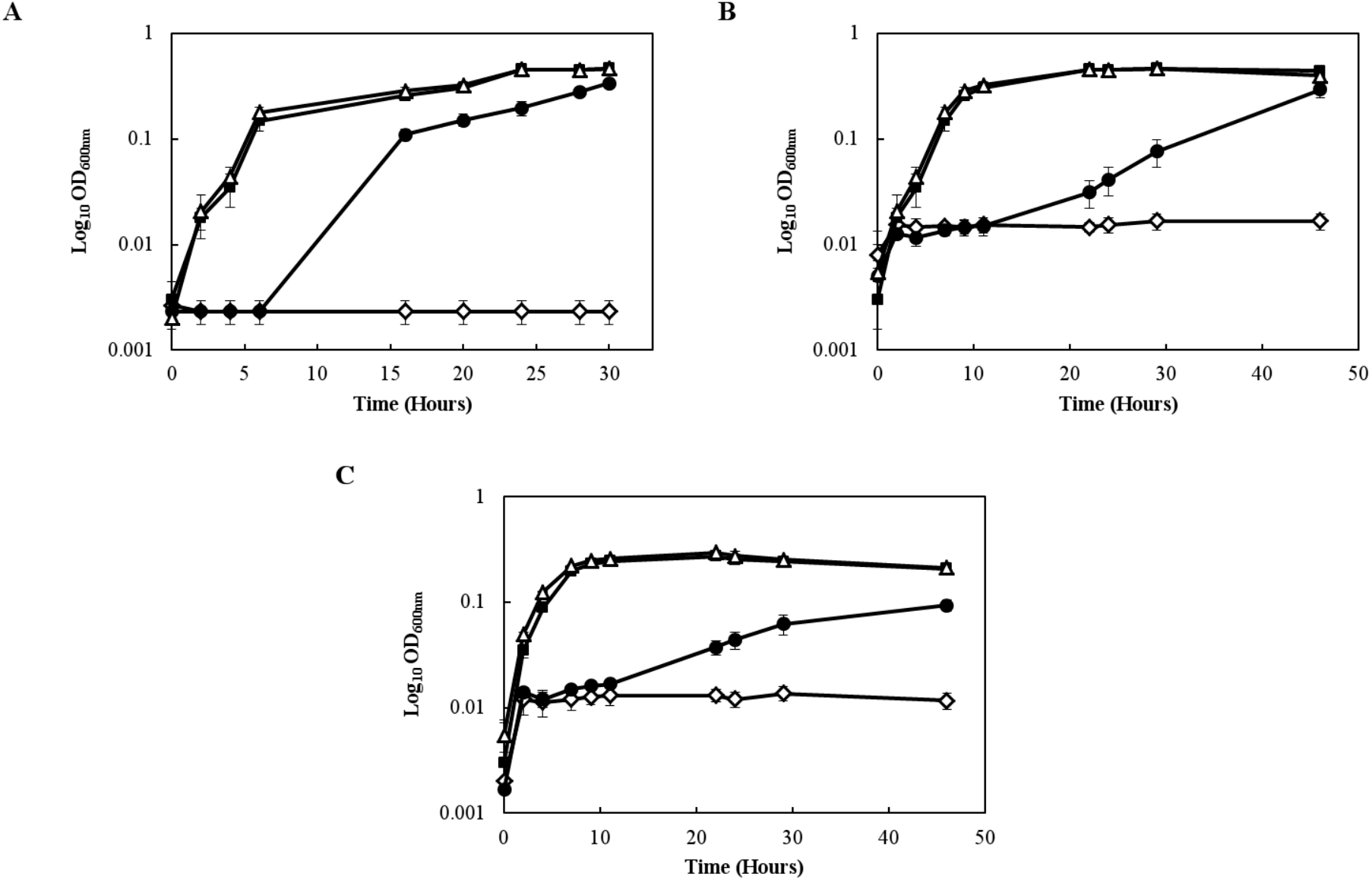
Quinol oxidase functionality of ImoA in CymA-dependent respiratory pathways. The ability of ImoA to reduce multiple periplasmic proteins was tested by quantifying growth of Δ*cymA* containing pBBR1MCS-2::*imoA* on different electron acceptors including fumarate (A), DMSO (B) and nitrate (C) while using lactate as the electron donor. *S. oneidensis* containing pBBR1MCS-2 (Δ), Δ*cymA* containing the empty pBBR1MCS-2 vector (◊), Δ*cymA* containing pBBR1MCS-2::*imoA* (●) and Δ*cymA* containing pBBR1MCS-2::*cymA* (■). Error bars represent standard deviation of the mean from experiments performed in triplicate.

### Efficient reduction of ferric citrate by ImoA requires both STC and FccA

To evaluate the ability of ImoA to exchange electrons with STC and FccA, the two periplasmic proteins reported to shuttle electrons from CymA to MtrA in *S. oneidensis*, strains missing CymA and STC (Δ*cymA* Δ*cctA*), CymA and FccA (Δ*cymA* Δ*fccA*) and all three (Δ*cymA* Δ*cctA* Δ*fccA*) were used. ImoA was expressed in these backgrounds and the strains were evaluated for the reduction of ferric citrate (**Figure 6**). The data show that ImoA is able to exchange electrons with both STC and FccA in *S. oneidensis*. However the reduction of ferric citrate only occurred after approximately 10 hours, compared to a wild-type strain carrying the empty plasmid (**Figure 6**). Interestingly, some ferric citrate reduction activity was observed in the Δ*cymA* Δ*cctA* Δ*fccA* strain containing pBBR1MCS-2::*imoA* (**Figure 6**), suggesting that ImoA may be able to exchange electrons with periplasmic proteins other than FccA and STC for the reduction of ferric citrate.

**Figure 6.**
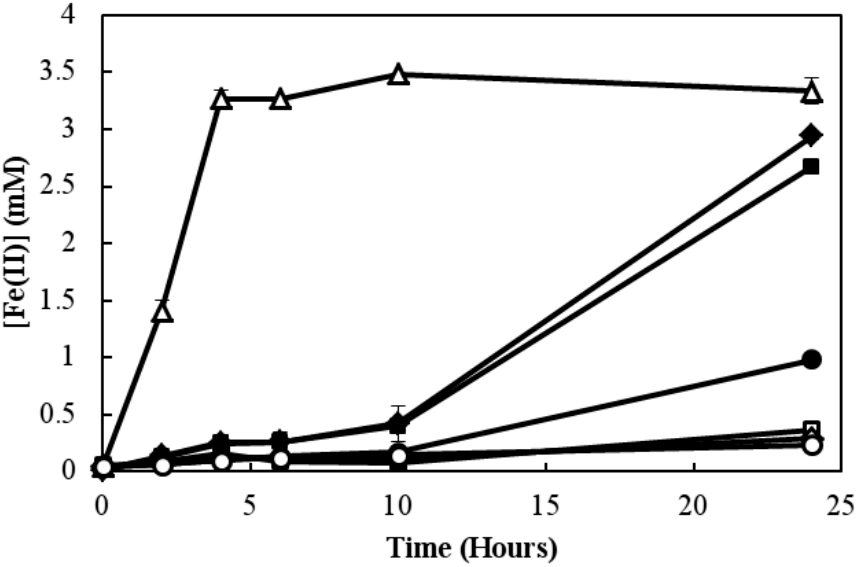
Ability of ImoA to exchange electrons with STC and FccA in *S. oneidensis*. Resting cell ferric citrate reduction by *S. oneidensis* containing pBBR1MCS-2 (Δ), Δ*cymA* Δ*fccA* containing pBBR1MCS-2 (◊), Δ*cymA* Δ*cctA* containing pBBR1MCS-2 (□), Δ*cymA* Δ*fccA* Δ*cctA* containing pBBR1MCS-2 (○), Δ*cymA* Δ*fccA* containing pBBR1MCS-2::*imoA* (◆), Δ*cymA* Δ*cctA* containing pBBR1MCS-2::*imoA* (■), Δ*cymA* Δ*fccA* Δ*cctA* containing pBBR1MCS-2::*imoA* (●). Error bars represent standard deviation of the mean from experiments performed in triplicate.

To further confirm the ability of ImoA to interact with STC and FccA we used NMR spectroscopy that is highly sensitive to changes in the chemical environment of nuclei, using purified proteins (17). By looking at the changes of the chemical shifts of the signals from the hemes of STC and FccA upon addition of ImoA (**Figure 7**), it is possible to observe that some of these signals change in position, indicating that an interaction between the proteins occurs. The assignment of these signals to specific hemes in the proteins STC and FccA already available in the literature (29, 30), provides the identification of the interaction site. While in FccA only signals from heme I are affected upon addition of ImoA (**Figure 7A**), in STC signals from the methyl groups of hemes II, III and IV are changed when ImoA is added (**Figure 7B**).

**Figure 7.**
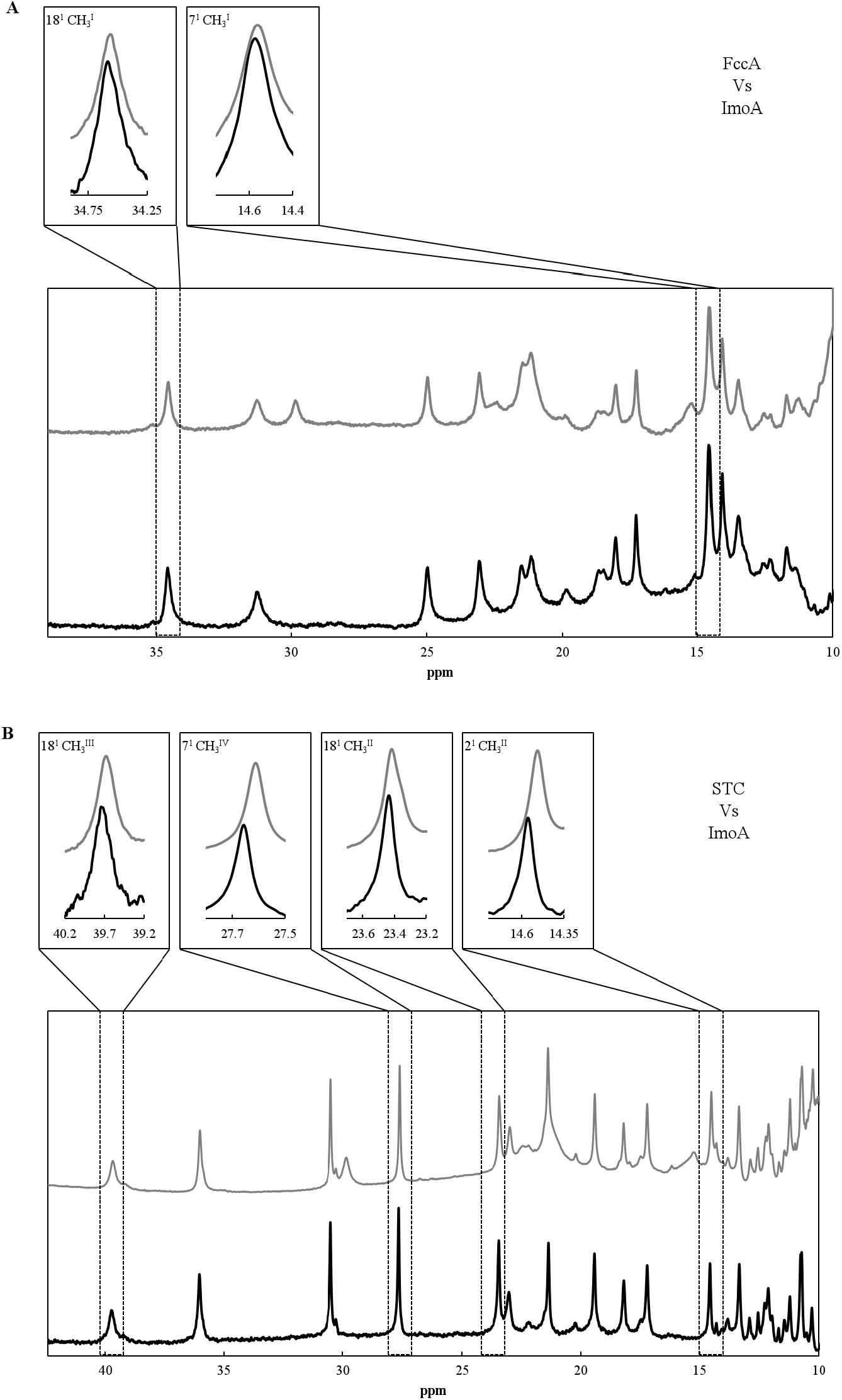
^1^H-1D NMR spectral changes of FccA (A) and STC (B) in the presence of ImoA. The bottom spectra was obtained with the individual proteins, while the top spectra was obtained upon addition of ImoA (molar ratio of [ImoA]/[STC or FccA] = 2). Boxes show examples of methyl signals with significant chemical shift perturbation. The methyl group is identified using the IUPAC-IUB nomenclature for hemes. The Roman numeral corresponds to the order of heme binding to the polypeptide chain.

## Discussion

Chemolithoautotrophic FeOB play an important role in the biogeochemical cycling of iron and other elements in numerous environments. However, the functionality of the electron transfer proteins required for their growth and survival in the environment remain poorly understood. The *c*-type cytochromes of the NapC/NirT family are components of electron transport pathways, and have been found to be encoded in the genomes of chemolithoautotrophic FeOB. In *S. lithotrophicus* ES-1 ImoA is a *c*-type tetraheme cytochrome of the NapC/NirT family proposed to act as a quinone reductase, receiving electrons from iron oxidizing extracellular electron uptake pathway to reduce the quinone pool (6). In this organism a *bc*_1_ complex that may act as a quinone reductase also exists, and it is unclear what, if any, advantages would be gained by having two quinone reductase systems. In order to understand the role of this protein in *S. lithotrophicus* ES-1, the biochemical, structural and functional properties of ImoA were investigated.

The characterization of ImoA showed that this protein is a monomer with one high-spin and three low-spin hemes (**Figure 2**). Phylogenetically, this protein is clustered closer to NirT sequences than to CymA from *Shewanella* sp. (**Figure 1**). Sequence alignment of the NapC/NirT family members (**Figure S4**) and the structure of NrfH enable us to propose the axial ligands of the hemes of ImoA. The CXXCH motifs required to bind the four *c*-type hemes (I-IV) are highly conserved in all these proteins, with the cysteines being vital to covalently bind the heme to the apo-protein, and the histidines being the proximal axial ligand of the hemes (31). The low-spin hemes are bis-His coordinated by conserved distal histidines with His186, His74 and His167, being most likely candidates for the distal axial ligands of hemes II, III and IV, respectively. The high-spin heme of ImoA, identified by RR spectroscopy, is suggested to be heme I, analogous with NrfH and CymA proteins of the NapC/NirT family (32, 33). In *Desulfovibrio vulgaris* heme I from NrfH is ligated by a methionine residue, Met49 of the CXXCHXM motif, as the proximal axial ligand instead of the histidine of the CXXCH heme *c*-binding motif (33). Although this methionine residue is completely conserved in NrfH proteins, it is only present in some sequences of NapC (34), and is absent in CymA (35). ImoA also contains one methionine after the heme *c*-binding motif of heme I, the Met61 (**Figure S4**). However, the NMR spectrum of reduced ImoA in the presence of a strong-field ligand (e.g. azide and cyanide) to achieve octahedral coordination of the heme did not show any signal corresponding to a methionine axially coordinated to the heme (around −3 ppm) (36, 37). These experimental data suggest that in ImoA the histidine from the CXXCH motif is the proximal axial ligand of heme I, as is typical of *c*-type hemes. Although mutagenesis studies demonstrated that the methionine is essential for menaquinol oxidation in *Wolinella succinogenes* NrfH (38), it may not be crucial for the oxidation of the quinol pool, since in several quinone interacting proteins this residue is not present (34, 35). Interestingly, the lysine residue (Lys91 in CymA from *Shewanella* sp. strain ANA-3, Lys103 in NapC from *E. coli* and Lys82 in NrfH from *D. vulgaris*) predicted to be important for menaquinol interaction, electron transfer and possible proton translocation (39) is present in ImoA (as Lys101). Overall, ImoA contains several structural features that are considered to be crucial for menaquinol oxidation (i.e. high-spin heme I and Lys101), suggesting that it can function as a quinol oxidase.

*S. lithotrophicus* ES-1 has been reported to be metabolically limited, and only known to grow using either Fe(II) or thiosulfate oxidation (23). Given this, we used *S. oneidensis* as a surrogate to study the ability of ImoA to replace CymA under different respiratory conditions and obtain insights into the function of ImoA. While the reduction of iron citrate and electrodes by *S. oneidensis* containing ImoA were very similar to that of strains containing CymA (**Figure 4**), the reduction of other electron acceptors were less efficient with ImoA (**Figure 5**). This may be explained by the differences in interaction between CymA and ImoA with their physiological partners. Both STC and FccA are known to interact with CymA (17, 22). STC and FccA are both capable of transferring electrons across the periplasm and are required for the reduction of solid and soluble electron acceptors, including fumarate, iron citrate, electrodes and DMSO (18). ImoA appears to interact with both STC and FccA *in vivo*, as it can replace CymA in the double mutants Δ*cymA*Δ*cctA* and Δ*cymA*Δ*fccA* strains during the reduction of iron citrate (**Figure 6**). However, while CymA interacts with STC mainly by heme IV (17), the interaction of ImoA with STC affects most of the hemes (hemes II, III and IV). Furthermore, the interaction of ImoA and CymA with FccA are also distinct. ImoA interacts in the vicinity of heme I of FccA, while CymA interacts in the vicinity of heme II (17)). These differences may explain the slower reduction of the different substrates by the knock-out strains containing ImoA compared to CymA. Although ImoA is redox active in the same potential range as CymA (i.e. between 0 and −400 mV vs SHE, **Figure 3**) (32, 40), the reduction potential of the individual hemes may differ, which can lead to a distinct electron transfer mechanism (41). However, to investigate this, it will be necessary to discriminate each of the hemes in the protein, and evaluate their redox behavior individually, which for membrane bound multiheme cytochromes such as CymA and ImoA is still highly challenging (42, 43). It is tempting to speculate that the high spin heme that we tentatively identify as heme I in ImoA, may play an important role as an immediate electron donor (or acceptor). It is noteworthy that the redox potential of ImoA is adequate to extract electrons from low-potential quinone pools such as menaquinones, which have been shown to be the main substrate for CymA (44). While aerobic electron transport chains are typically dependent on ubiquinone, anaerobic electron transport chains are typically dependent on menaquinone (45). Although ImoA is able to oxidize menaquinol in *S. oneidensis*, genes encoding menaquinone biosynthesis are absent from the *S. lithotrophicus* ES-1 genome. As expected, a complete pathway for ubiquinone biosynthesis is present in the *S. lithotrophicus* ES-1 genome.

The potential role of ImoA as a quinol oxidase in *S. lithotrophicus* ES-1 is intriguing from a physiological and ecological perspective. Fe(II) and thiosulfate oxidation pathways are dependent on electron transport chains that require quinone reductases rather than a quinol oxidase (23). Oxidoreductases from NapC/NirT family can be found in both Fe(III)-reducing bacteria and in *S. lithotrophicus* ES-1, which suggests that these NapC/NirT family proteins may catalyze both quinol-oxidizing and quinone-reducing reactions (13). If ImoA is a quinol oxidase, as indicated by our data, what role could it be playing in *S. lithotrophicus* ES-1? By analyzing the *S. lithotrophicus* ES-1 genome, we identified genes that encode putative periplasmic redox proteins that are known to either directly or indirectly accept electrons from NapC/NirT family quinol oxidases including *nirS* (Slit_1129) and several homologs of *nirM* (**Table S1**). The protein sequence encoded by *Slit_1129* is 68% identical to NirS from *P. aeruginosa*. NirM is known to transfer electrons from NirT to NirS during nitrite reduction (46). Interestingly, MtoD is homologous to NirM (28% identity and 43% similarity of respective protein sequences), suggesting that this pathway may also exist in *S. lithotrophicus* ES-1, although this bacterium has not been tested to reduce nitrogen compounds (23). Recently, *Sideroyxydans* sp. were found to be highly abundant on a denitrifying biocathode in a microbial electrochemical system containing arsenite (47), however its potential role in the redox cycle of arsenic or nitrogen compounds has not been studied. Further studies are still needed to fully understand how *S. lithotrophicus* ES-1 couples the oxidation of iron to the generation of NADH and a proton motive force. The insights obtained in this work will be extremely relevant for understanding the bioenergetic pathways of FeOB, allowing us to unravel their role in the biogeochemical cycling of iron, and their application in bioelectrosynthesis.

## Materials and Methods

### 1. Phylogenetic analysis

Amino acid sequences representing CymA, NapC, NirT, NrfH, cytochrome c552 and NapC/NirT proteins from *S. lithotrophicus* ES-1 and *Sideroxydans spp*. CL21 were downloaded from NCBI Data bases and aligned using ClustalW (48). MEGA7 was used to generate a phylogenetic tree using the maximum likelihood method with 2,000 bootstrap replications, based on the JTT matrix-based model (49, 50).

### 2. Construction of bacterial strains

Strains, primers and plasmids used in this study are listed in Table 1. The gene *imoA* was synthesized by NZYTech, amplified using the primers 1 and 2 presented in Table 1, and cloned in the NcoI restriction site of the pBAD202/D-TOPO (Invitrogen, Carlsbad, CA, USA) using the NEBuilder Assembly kit (New England BioLabs). To facilitate the purification process a Strep-tag sequence was included in the reverse primer (primer 2) before the stop codon of the gene. The resulting plasmid (pBAD202::*imoA*) was transformed into a chemically competent *E. coli* strain JM109(DE3) co-transformed with vector pEC86, which contains the *ccmABCDEFGH* genes (51).

**Table 1.**
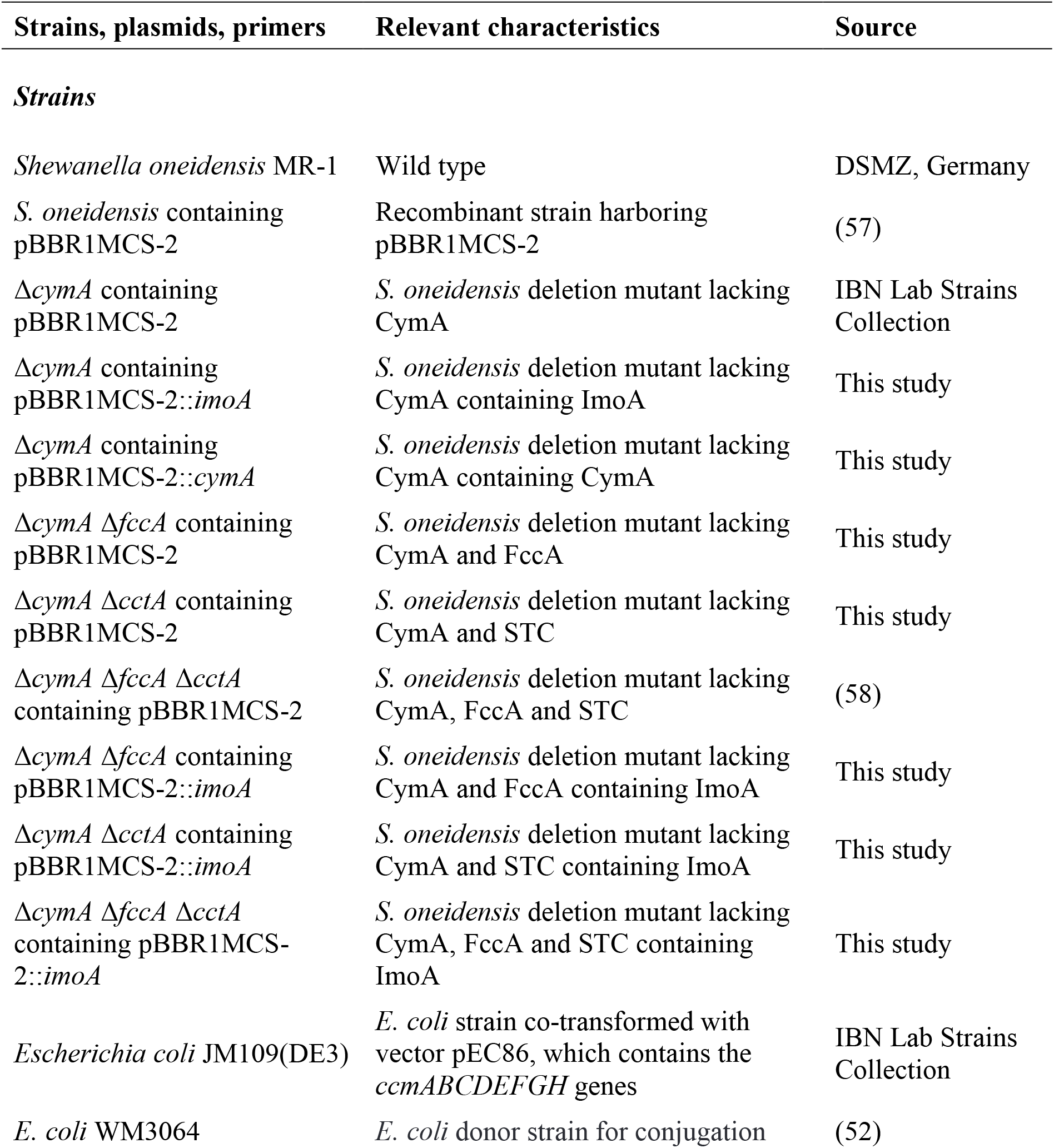
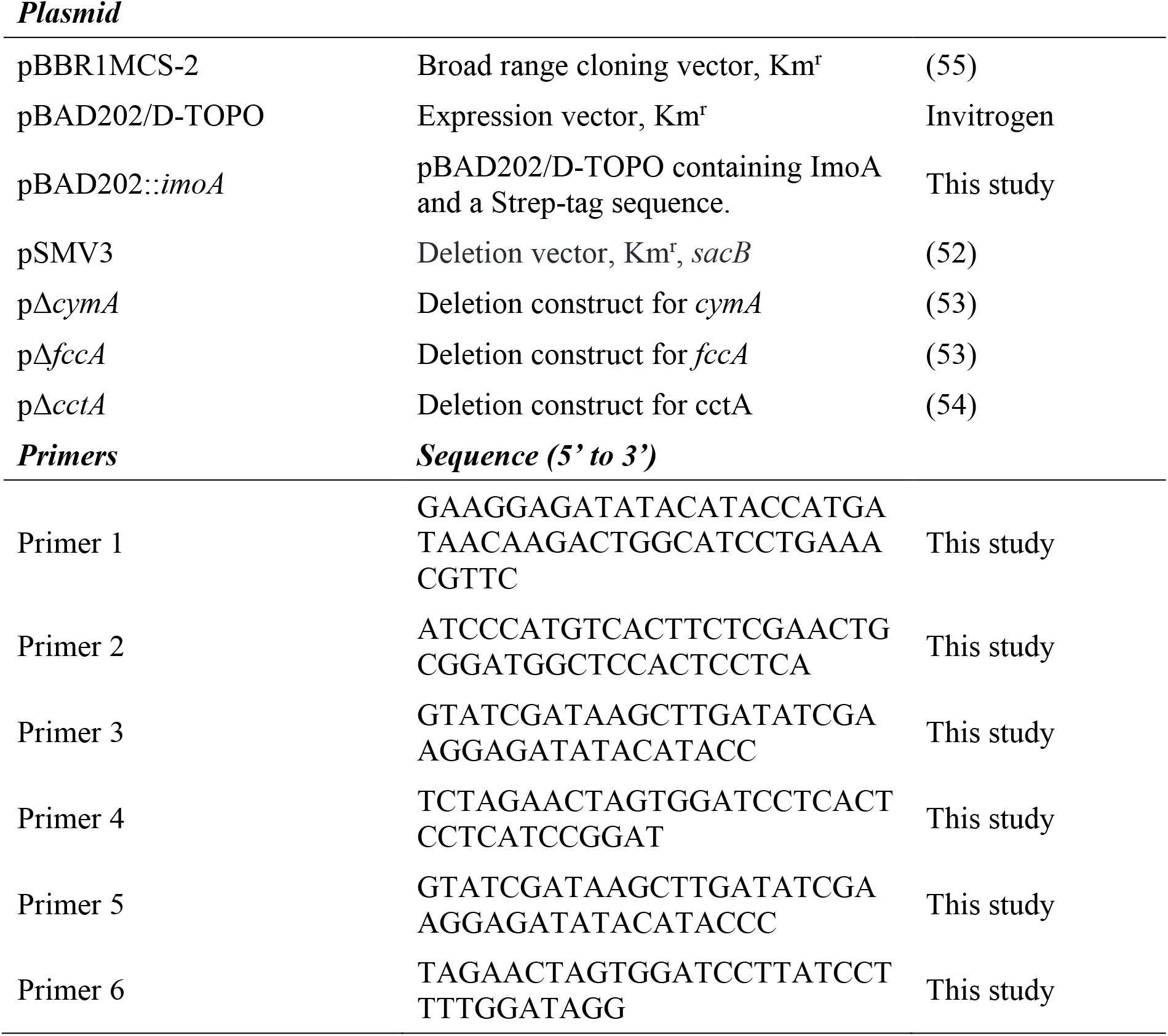
Bacterial strains, plasmids, and primers used in this study.

Deletion construct plasmids were generated by amplifying approximately 1 kb DNA fragments upstream and downstream of the targeted sequence, including nine nucleotides after the start codon and nine nucleotides before the stop codon, followed by ligating and cloning into the suicide vector pSMV3 (52) which was transformed into chemically competent *E. coli* WM3064 cells (53, 54). Transformed *E. coli* WM3064 cells were selected on Lysogeny Broth (LB) medium plates containing 50 μM kanamycin and 360 μM diaminopimelic acid (DAP) and used to conjugate deletion constructs into desired *Shewanella* strains. Merodiploids were selected by growth on LB plates containing 50 μM kanamycin without DAP. Merodiploids were resolved by sucrose counterselection mediated by SacB and screened by PCR.

For the studies of growth phenotype, the *imoA* and *cymA* genes were amplified using primers 3 and 4, and 5 and 6, respectively (Table 1), and cloned in the EcoRV and BamHI restriction site of the pBBR1MCS-2 (55) using the NEBuilder Assembly kit (New England BioLabs). pBBR1MCS-2 was a gift from Kenneth Peterson (Addgene plasmid # 85168; http://n2t.net/addgene:85168; RRID:Addgene_85168). The resulting plasmids were then transformed into *Shewanella* using electroporation (56). In these constructs, the ribosome binding site (RBS) present at the pBAD2020/D-TOPO vector was also included and the reverse primer contained a stop codon to guarantee that a native protein (i.e. without a tag) is expressed.

### 3. Protein characterization

#### 3.1. Production of ImoA

The *E. coli* strains JM109(DE3) harboring pEC86 and pBAD202::*imoA* were grown aerobically at 37 °C in Terrific Broth medium containing 50 μg/mL kanamycin and 34 μg/mL chloramphenicol in 5 L Erlenmeyer flasks containing 2 L of medium at 150 rev./min. Protein expression was induced by addition of 3 mM L-arabinose at optical density (OD_600nm_) of 1.0 and the cells were allowed to grow under the same conditions for 48 h before harvesting. Bacterial cells were harvested by centrifugation at 10,000 *g* for 15 min at 4 °C. The cell pellet collected from 6 L growth was resuspended in 20 mM HEPES (pH 7.8) containing protease inhibitor cocktail (Roche) and DNase I (Sigma). The disruption of the cells was achieved by three passages through a French Press at a pressure of 1000 psi (6.89 MPa). The crude extract was centrifuged at 200,000 g for 1 h at 4 °C (Beckman Coulter Optima LE-80K). The membrane pellet was homogenized and solubilized in 20 mM HEPES (pH 7.8) containing a protease inhibitor cocktail and 4 % (w/v) DDM at 4 °C overnight to solubilize the membrane proteins. The insoluble material was removed by ultracentrifugation at 200,000 *g* for 1 h at 4 °C. Harvested clear cell lysates containing the solubilized membrane proteins were loaded into a 5 mL Strep-Tactin® column (IBA) and the protein was eluted using Elution Buffer (20 mM HEPES, pH 7.8, 100 mM NaCl, 0.05 % DDM, 2.5 mM desthiobiotin). Eluted fractions were analyzed by SDS-PAGE (12 % gel) stained for heme proteins (59) and by UV-visible spectroscopy to select those containing pure protein. The purity index of the sample was defined by the A_Soret peak_/A_280nm_ ratio. To provide direct insight into the oligomeric state of ImoA, the sample was also concentrated and subjected to size-exclusion chromatography on a Superdex 200 10/300 GL column (GE Healthcare) in 20 mM HEPES, pH 7.8, 100 mM NaCl containing 0.05 % DDM. The identity of ImoA was confirmed by N-terminal sequencing and mass spectrometry and protein concentration was estimated using the absorption coefficient, ε_409nm_, of 125,000 M^−1^cm^−1^ per heme for the oxidized state of the protein (60).

#### 3.2. Spectroscopic Techniques

##### 3.2.1. UV-Visible Spectroscopy

UV-visible spectra of ImoA were acquired in 20 mM HEPES buffer (7.8) with 100 mM KCl and 0.05 % DDM on a Shimadzu UV-1800 spectrophotometer (Shimadzu, Gamby, OR, USA) in the 250 to 800 nm wavelength range at room temperature. Spectra in the reduced state were obtained by the addition of small volumes of concentrated sodium dithionite solution (Sigma) to the oxidized sample.

##### 3.2.2. NMR Spectroscopy

NMR experiments were performed at 25 °C on an NEO 500 MHz NMR spectrometer (Bruker, Rheinstetten, Germany) equipped with a 5 mm TCI C/N Prodigy Cryo probe. 10 % (v/v) of ^2^H_2_O (99.9 atom %) was added to the protein sample prepared solubilized in 20 mM HEPES buffer (pH 7.8) with 100 mM KCl and 0.05 % DDM before spectral acquisition. The ^1^H-1D-NMR spectrum was acquired with spectral width of 80 kHz, and processed in the Topspin 4.1.1 software from Bruker using an exponential apodization function. For solvent suppression and enhancement of the paramagnetic signals, a SuperWEFT pulse sequence was applied. Chemical shifts are reported in parts per million (ppm) and the proton spectrum was calibrated using the water signal as the internal reference.

##### 3.2.3. RR Spectroscopy

RR spectra were acquired using a confocal Raman spectrometer (Jobin-Yvon LabRam 800 HR, HORIBA) equipped with a liquid nitrogen-cooled CCD detector, employing 406 nm excitation from a krypton ion laser (Coherent INNOVA 300c). Spectra were measured from 100 μL of 100 μM of ImoA placed into a rotating cuvette (Hellma) to prevent prolonged exposure of individual protein molecules to laser irradiation. Spectra were recorded with 0.8 mW laser power and 20 s accumulation time at room temperature; 8 spectra were co-added in order to improve signal-to-noise ratio. Spectral deconvolution was performed using LabSpec 5.4 software.

#### 3.3. Cyclic Voltammetry

Protein film voltammetry was carried out at 25 °C using a three-electrode electrochemical cell configuration consisting of a PGE working electrode (IJ Cambria Scientific), an Ag/AgCl (3 M KCl) reference electrode and a graphite rod counter electrode. The experiments were performed inside a Coy anaerobic chamber using an electrochemical analyser from CHI Instruments controlled by the manufacturer’s software (version 20.04). The PGE electrode was cleaned and freshly polished before every experiment. For these experiments 5 μl of pure protein of ImoA solubilized in 20 mM HEPES buffer (pH 7.8) with 100 mM KCl and 0.05 % DDM (100 μM) was deposited on the surface of the working electrode and left to dry for approximately 30 min. Excess and/or unattached protein was removed by rinsing the electrode with distilled water. The electrode was then immersed in 3 mL of a mixed buffer solution containing 5 mM of HEPES (Sigma Aldrich), 5 mM of MES (Sigma Aldrich) and 5 mM of TAPS (Sigma Aldrich) and 100 mM KCl. The desired pH values were adjusted with 1 M NaOH. Voltammograms were acquired at a scan rate of 0.4 Vs^−1^. After each experiment, the solution was removed, and the pH measured. The baseline was obtained in similar conditions but without depositing the protein on the electrode. QSoas program (version 1.0) available at https://bip.cnrs.fr/groups/bip06/software/ (61) was used to subtract the capacitive current and suppress background noise of the raw electrochemical data. All potentials are reported with respect to a standard hydrogen electrode (SHE) by addition of 210 mV (62) to those measured.

### 4. Fe(III) citrate reduction assay

Fe(III) citrate reduction assays were performed as described in (63). Briefly, cells were freshly struck from −80 °C glycerol stocks to lysogeny broth (LB) plates containing 50 μg/mL kanamycin. Oxic LB liquid medium containing 50 μg/mL kanamycin were inoculated with single colonies and incubated in a shaker at 30 °C. The cells were washed with *Shewanella* basal medium (SBM) (57) and resuspended in the same medium to obtain a cell density of 10^9^ cells/mL. 30 μL of the resuspended cells were added to 270 μL of the SBM medium containing 20 mM sodium lactate and 5 mM of ferric citrate on a 96 well plate. The 96 well plate was placed inside an anaerobic chamber which was made anaerobic by flushing with oxygen free argon. The anaerobic chamber was incubated at 30 °C. Samples were collected periodically to quantify Fe(II), produced as a result of Fe(III) reduction, using the ferrozine assay (64).

### 5. Growth on anodes

To evaluate the ability of the Δ*cymA* strains carrying *imoA* or *cymA* (Table 1) to perform extracellular electron transfer to anodes, the strains were grown in three-electrode bioelectrochemical reactors. The reactors were single chambered and made out of 100 mL Schott glasses closed with butyl rubbers and fixed by screw caps to guarantee anaerobic conditions during the experiment. The working anode was made of graphite felt (GFD 2.5 from Sigracell (Germany), round size of 13 mm diameter) and stamped into a self-made electrode holder (hungate screw cap) with a silicon stopper. The counter electrode was a graphite rod and both electrodes were connected with titanium wires. The used Ag/AgCl (3 M KCl) reference electrode (IJ Cambria) and working and counter electrode were inserted through a hole previously drilled into the rubber. The surface of the anode electrode in contact with the medium was of 9 mm diameter (= 0.64 cm^2^). Prior to autoclaving, the working electrode was immersed in isopropanol and washed with deionized water. Before use, the reactors were filled with deionized water and autoclaved.

Seven bioelectrochemical experiments were conducted in parallel: one abiotic control, a triplicate of one bacterial strain and a triplicate of another bacterial strain. Δ*cymA* containing an empty pBBR1MCS-2 vector was used as the control strain. The bacteria were grown overnight in oxic LB medium containing 50 μg/mL kanamycin, washed with SBM and inoculated in anoxic SBM containing 20 mM sodium lactate in the reactors. All the experiments were performed at 30 °C using a Dropsens multipotentiostat in the chronoamperometric mode applying 200 mV vs. Ag/AgCl and measuring the current every 30 seconds. Approximately 2 hours after the experiment started, washed *Shewanella* cells were added to the reactors with a resulting OD_600nm_ of 1.0.

### 5. Growth quantification

For growth curve experiments, *S. oneidensis* strains containing the desired plasmid were freshly struck from −80 °C glycerol stocks to LB plates containing 50 μg/mL kanamycin. Oxic LB liquid medium containing 50 μg/mL kanamycin were inoculated with single colonies and incubated in a shaker at 30 °C. Overnight cultures were washed with SBM and inoculated into anoxic SBM medium containing 20 mM sodium lactate and 40 mM of the respective electron acceptors. Growth of different strains was measured over time using OD_600nm_.

### 6. NMR interaction studies

Stock samples of STC and FccA (17) in 20 mM HEPES (pH 7.8) with 100 mM KCl and 0.05 % DDM were lyophilized and dissolved in ^2^H_2_O (99.9 atom %). NMR spectra obtained before and after lyophilization were identical, demonstrating that the protein structure was not affected by this procedure (data not shown). Samples containing 50 μM of these proteins were titrated with ImoA solubilized in 20 mM HEPES (pH 7.8) with 100 mM KCl and 0.05 % DDM prepared in ^2^H_2_O (99.9 atom %) using a ratio of 1:2. ^1^H-1D-NMR spectra were recorded before and after the addition of ImoA. Given that the chemical shifts of the signals observed in the NMR spectra correspond to the heme methyl substituents of STC and FccA previously assigned to specific hemes in the protein structure (29, 30), any change in these signals upon the addition of ImoA indicates an interaction between the proteins, and allow the identification of the docking site in the *Shewanella* proteins.

## Acknowledgements

Financial support was provided by European EC Horizon2020 TIMB3 (Project 810856). This work was funded by national funds through FCT–Fundação para a Ciência e a Tecnologia, I.P. (FCT), ProjectMOSTMICRO-ITQB with refs UIDB/04612/2020 and UIDP/04612/2020, and projects PTDC/BIA-BQM/30176/2017 and PD/BD/135153/2017 integrated in the PhD Programme in NMR applied to chemistry, materials and biosciences (PD/00065/2013). The NMR data were acquired at CERMAX, ITQB-NOVA, Oeiras, Portugal with equipment funded by FCT, project AAC 01/SAICT/2016. N-terminal sequencing service was provided by the ITQB Research facilities, while spectrometry data was obtained by the MasSpectrometry Laboratory, Analytical Services Unit at ITQB/IBET. This work was also supported by the Office of Naval Research (N0014-21-1-2166) and the National Science Foundation (MCB-1815584).

